# Genomic architecture and sexually dimorphic expression underlying immunity in the red mason bee, *Osmia bicornis*

**DOI:** 10.1101/2021.10.04.462642

**Authors:** Jannik S. Möllmann, Thomas J. Colgan

**Author notes:** **Corresponding authors:** JSM, TJC.

## Abstract

Insect pollinators provide crucial ecosystem services yet face increasing environmental pressures. The challenges posed by novel and reemerging pathogens on bee health means we need to improve our understanding of the immune system, an important barrier to infections and disease. Despite its importance, for certain ecologically important species, such as solitary bees, our understanding of the genomic basis and molecular mechanisms underlying immune potential, and how intrinsic and extrinsic factors may influence immune gene expression is lacking. Here, to improve our understanding of the genomic architecture underlying immunity of a key solitary bee pollinator, we characterised putative immune genes of the red mason bee, *Osmia bicornis*. In addition, we used publicly available RNA-seq datasets to determine how sexes differ in immune gene expression and splicing but also how pesticide exposure may affect immune gene expression in females. Through comparative genomics, we reveal an evolutionary conserved set of more than 500 putative immune-related genes. We found genome-wide patterns of sex-biased gene expression, including immune genes involved in antiviral-defence. Interestingly, the expression of certain immune genes were also affected by exposure to common neonicotinoids, particularly genes related to haemocyte proliferation. Collectively, our study provides important insights into the gene repertoire, regulation and expression differences in the sexes of *O. bicornis*, as well as providing additional support for how neonicotinoids can affect immune gene expression, which may affect the capacity of solitary bees to respond to pathogenic threats.

## Introduction

Insect pollinators provide key ecosystem services that are essential for the maintenance of agricultural crop yields, as well as natural biodiversity (Klein, Steffan-Dewenter and Tscharntke, 2003; Losey and Vaughan, 2006; Gallai *et al.*, 2009). Pollination by insects, including social and solitary bees, is estimated to contribute $15.2 billion to the US economy demonstrating the economic benefits provided by such services (Calderone, 2012). Despite the importance of such services, recent documented declines in bee populations have raised concerns over the continued provision of such services and related issues with food security (Vanengelsdorp and Meixner, 2010; *European Red List of Bees*, 2014; Goulson *et al.*, 2015). Both abiotic and biotic factors have been highlighted as contributing factors to decline, including habitat loss and fragmentation, climate change, increased pesticide usage in modern agriculture, as well as pathogens and disease (Brown and Paxton, 2009; Goulson *et al.*, 2015).

An important barrier to infection and establishment of disease is the invertebrate immune system (Rolff and Reynolds, 2009; Sadd and Schmid-Hempel, 2009). Despite lacking the adaptive immune system found in vertebrates, invertebrates have a dynamic innate immune system consisting of recognition molecules, signalling pathways and effector molecules, which coordinate the targeting and removal of potentially harmful entities (Hoffmann, 1995). In addition to the evolutionary importance of the immune system (Sadd and Schmid-Hempel, 2008; Viljakainen, 2015), understanding insect immunity has applied purposes, especially within the fields of biomedical, agricultural and conservation biology. Genomic studies on insects have provided novel insights into the genes and genomic architecture underlying the immune system (Christophides *et al.*, 2002; Evans *et al.*, 2006; Sackton *et al.*, 2007; Waterhouse *et al.*, 2007; Gerardo *et al.*, 2010). Such studies have documented and helped understand the immune potential and capacity of a species through the identification of gene family expansions, contractions, as well as lineage-specific or novel genes that demonstrate immune function (Adams, 2000; Evans *et al.*, 2006; Barribeau *et al.*, 2015). Indeed, comparative genomics allows for examining the types of selection pressures acting on such genes providing important insights into their evolutionary history. Given the enormous selection pressures placed upon hosts by pathogens (Combes, 2001), genes involved in the immune system are expected and have been observed to evolve under strong positive selection. Indeed, in comparative genomic studies of both vertebrates and invertebrates, including insects, immune genes are often identified with signatures of accelerated rates of evolution (Viljakainen *et al.*, 2009; Roux *et al.*, 2014; Shultz and Sackton, 2019).

Functional genomics, such as genome-wide transcriptional profiling (“RNA-seq”), provide high-scale resolution of genome-wide changes in gene expression in response to immune or pathogen challenge but also intrinsic differences in expression between different life cycle stages or sexes (Fish, 2008; Klein and Flanagan, 2016). Indeed, sexually dimorphic gene expression has been identified across taxonomically diverse groups and may exist due to differences in life histories, hormonal abundance, biochemical reactions or sex-specific genomic architecture (Hill-Burns and Clark, 2009). Such differences can underlie differences in immune expression and function but also susceptibility to disease and related survival (Ingersoll, 2017). A striking example whereby sex-specific differences in genomic architecture may underlie differences in immune potential, activity and expression are members of the Hymenoptera, which include the bees, ants and wasps. Within this group, sexes differ in their ploidy with females developing from fertilised diploid eggs while males develop from unfertilised haploid eggs (Pamilo and Crozier, 1981). The haploid nature of males means that any alleles carried are automatically expressed and open to selection, which can result in the rapid removal of maladaptive deleterious alleles from the gene pool (Joseph and Kirkpatrick, 2004). The haploid nature of males has led to predictions that they are more susceptible to environmental challenges, such as pathogens (O’Donnell and Beshers, 2004), although empirical evidence to support this view has been conflicting (Baer *et al.*, 2005; Calleri *et al.*, 2006; Ruiz-González and Brown, 2006; Colgan *et al.*, 2011; Retschnig *et al.*, 2014).

Despite the fact that pathogen exposure and intrinsic differences can affect or influence immune gene expression, other factors can also have an influence, including nutritional status (Moret and Schmid-Hempel, 2000; DeGrandi-Hoffman and Chen, 2015), mating (Peng, Zipperlen and Kubli, 2005; Lawniczak *et al.*, 2007; Barribeau and Schmid-Hempel, 2017), periods of senescence or dormancy (Nakamura *et al.*, 2011; Kubrak *et al.*, 2014; Colgan *et al.*, 2019), as well as environmental factors, such as temperature (Xu and James, 2012; Chen, Nolte and Schlötterer, 2015). One environmental challenge that has received considerable attention of late is the influence of pesticide exposure on immune expression and function. Chemical pesticides, such as pyrethroids and neonicotinoids, interact with the insect nervous system resulting in paralysis and death (Matsuda *et al.*, 2001). The efficacy of chemicals, such as neonicotinoids, combined with its lower toxicity to vertebrates, their systemic mode of action, as well as the lack of requirement for reapplication has resulted in their increased use in modern agricultural practices (Jeschke and Nauen, 2008). Despite their efficacy in killing agricultural pest species, the ubiquitous distribution of neonicotinoids across tissues means that non-target insects, including beneficial pollinators, may come in contact with such chemicals through food resources, such as nectar and pollen (Blacquière *et al.*, 2012). The concentration of such chemicals can result in sublethal and indirect effects on the insect phenotype and has been identified to adversely affect the behaviour (Gill, Ramos-Rodriguez and Raine, 2012; Stanley, Smith and Raine, 2015; Arce *et al.*, 2017, 2018; Siviter *et al.*, 2018), neurobiology (Moffat *et al.*, 2015, 2016) and gene expression of key ecological and commercial pollinators (Chaimanee *et al.*, 2016; Beadle *et al.*, 2019; Bebane *et al.*, 2019; Colgan *et al.*, 2019). In addition, such chemicals have been shown to affect immune expression and function (Di Prisco *et al.*, 2013; Mason *et al.*, 2013; Chmiel *et al.*, 2019; Brandt *et al.*, 2020) raising concerns that these effects may influence the ability of an exposed individual to respond to pathogenic threats (James and Xu, 2012). While studies have identified molecular mechanisms by which bees can metabolise certain neonicotinoids (Manjon *et al.*, 2018; Beadle *et al.*, 2019; Troczka *et al.*, 2019), our understanding of changes in immune expression due to neonicotinoid exposure, especially for solitary bees, is limited.

The mason bees (*Osmia* species) are an important group of solitary bee pollinators but are generally understudied from an immunological perspective. One such species is the red mason bee *Osmia bicornis* (Order Hymenoptera; Family Megachilidae), a common pollinator found across central Europe, which has been increasingly incorporated into modern agricultural practices (Gruber *et al.*, 2011). Despite its importance, it faces a number of environmental challenges that can influence the immune system, including pathogens and parasites (Seidelmann, 2006; Schoonvaere *et al.*, 2018; Tian *et al.*, 2018; Bramke *et al.*, 2019), as well as pesticides (Brandt *et al.*, 2020). Similarly, intrinsic differences between the sexes, which differ in morphology, physiology, behaviour and ploidy (Dmochowska-Ślęzak *et al.*, 2015; Rogers, Frasnelli and Versace, 2016; Szentgyörgyi *et al.*, 2017), may result in differences in immune expression and associated susceptibility to pathogenic threats. However, at present, our understanding of the immune gene repertoire and expression in *O. bicornis* is currently limited.

To improve our understanding on the immune potential of *O. bicornis*, we performed a comparative genomic analysis to identify the immune gene repertoire of the red mason bee as well as identify potential contractions and expansions of important gene families and determine whether the red mason bee is missing immune genes. Furthermore, to understand how genes underlying immunity may be expressed differently between the sexes, we investigated evidence of sex-biased gene expression and alternative splicing. Lastly, as pesticides can negatively affect different aspects of *Osmia* health, including developmental rate (Mokkapati, Bednarska and Laskowski, 2021), foraging behaviour (Boff *et al.*, 2021; Straub *et al.*, 2021), reproductive output (Sandrock *et al.*, 2014; Woodcock *et al.*, 2017; Ruddle *et al.*, 2018), thermoregulation (Azpiazu *et al.*, 2019), as well as impact immune function in red mason bees (Brandt *et al.*, 2020), we examined whether neonicotinoid-exposed individuals differed in immune gene expression.

## Results

### Putative expanded immune gene repertoire in the Hymenoptera

To infer putative immune genes in *O. bicornis*, we independently examined the presence of homologues of previously characterised immune genes of the fruit fly, *Drosophila melanogaster*, and the closely related earth bumblebee, *Bombus terrestris* in the *O. bicornis* predicted proteome. We merged resulting *O. bicornis* homologues of known immune genes in *D. melanogaster* and *B. terrestris*, resulting in a total of 541 putative immune gene homologues, of which 99 were only found among the set of *B. terrestris* immune genes, 387 were only found among the set of *D. melanogaster* immune genes and 55 were found among both sets (Figure 1A, Supplemental File S1). We also ran functional enrichment analyses and found enrichment in immune terms for the putative immune genes derived uniquely via homology to *D. melanogaster* and for the putative immune genes found among both sets while for the putative immune genes derived uniquely via homology to *B. terrestris* we found enrichment in terms only indirectly linked to the immune system such as “response to stress” and “response to toxic substance” (Supplemental File S2).

**Figure 1.**
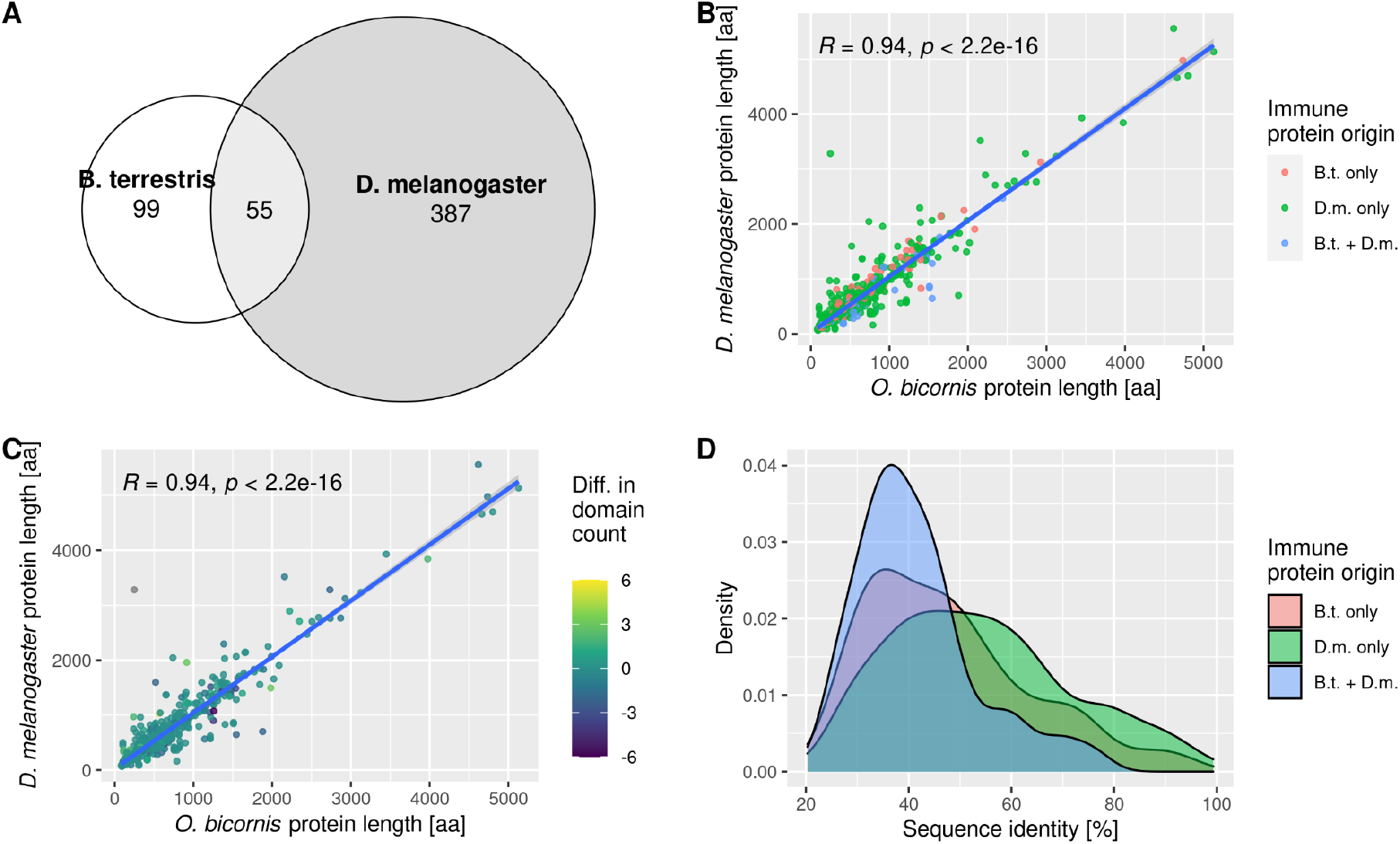
Expanded repertoire of putative immune genes in the red mason bee, *Osmia bicornis*. **(A)** Overlap of putative *O. bicornis* immune genes inferred via homology to *B. terrestris* and *D. melanogaster*. **(B-C)** Correlation of protein lengths (amino acids) and number of functional domains shared between *O. bicornis* and *D. melanogaster* protein homologues with colours indicating **(B)** immune gene set where homology was found (“B.t. only” = homologues identified through comparison with *B. terrestris*; “D.m. only” = homologues identified through comparison with *D. melanogaster*; “B.t.+D.m.” = homologues identified in comparisons with both species) and **(C)** difference in number of domains (yellow: higher domain count in *D. melanogaster*, purple: higher domain count in *O. bicornis)*. **(D)** Distribution of sequence identity (percentage of identical amino acid positions) of homologous protein pairs between *D. melanogaster* and *O. bicornis* for putative *O. bicornis* immune proteins. Colours indicate the species through which homologues were identified.

Given that previous genomic studies on immune gene repertoires in hymenopterans have described lower gene counts, and since *O. bicornis* is more evolutionarily distantly related to *D. melanogaster* than *B. terrestris*, we further examined immune genes identified only through homology to *D. melanogaster* to determine confidence in homology. For this, we examined the sequence identity of homologous immune proteins as calculated by OrthoFinder between *O. bicornis* and *D. melanogaster* revealing higher percentage sequence identity for putative immune proteins found only in the set of *D. melanogaster* immune proteins compared to immune proteins found uniquely in the set of canonical *B*.*terrestris* immune proteins or immune proteins found in both sets (Figure 1D). As metrics of sequence identity can be influenced by protein length, we compared predicted protein lengths between homologous pairs, revealing a strong positive correlation (Pearson’s Product Moment Correlation Coefficient, R=0.94, *p*< 2.2e-16, Figure 1B-C). Lastly, we examined if homologous sequences shared the same type and number of functional domains, which would potentially suggest conserved function. We found identical domain annotations for a high percentage (71.01%, n=722) of all homologous pairs between *O. bicornis* and *D. melanogaster* with on average 90.05% of the domains shared between pairs.

Our analysis also revealed immune genes potentially missing in *O. bicornis*. We did not identify homologues for 232 *D. melanogaster* immune genes as well as one canonical *B. terrestris* immune gene, the antimicrobial peptide abaecin, in *O. bicornis* or its sister taxa, *O. lignaria* (Supplemental File S1). Inversely, as *Osmia* may contain lineage specific genes, including genes with potential immune functions, we identified genes (n=78, split across 48 orthogroups; Supplemental File S1) shared between *O. bicornis* and *O. lignaria* that lacked homologues in the predicted proteomes of all other 19 insect species we used to infer homology relations (for species list see Experimental Procedures). Among the *Osmia*-specific genes that were annotated with at least one domain (n=24), we find 16 genes annotated with ribonuclease-domains (IPR036397, IPR012337) as well as one gene (LOC114879997) annotated with a rhabdovirus nucleoprotein domain (IPR004902).

To understand variation in the immune gene complement of the red mason bee, we compared the number of putative *O. bicornis* immune genes in conserved gene families and pathways to the number of homologues in three closely-related hymenopteran species (*O. lignaria, B. terrestris* and *A. mellifera*) and one more distantly-related fly species (*D. melanogaster*). We found similar numbers of genes in the four hymenopterans for most signaling pathways or non-pathway gene families, with slightly lower gene numbers in the two *Osmia* species compared to the other two hymenopterans for the Immune deficiency (ImD) and Toll pathways (Figure 2A) and higher numbers for inhibitors of apoptosis in *B. terrestris* (n=13) compared to the other hymenopterans (n=4 to 5, Figure 2B). When comparing the hymenopterans to *D. melanogaster*, the latter has higher gene counts for six out of 14 signaling pathways and 13 out of 22 non-pathway gene families but similar numbers for all other pathways and gene families. In addition, we checked the number of homologues in *O. bicornis* for *D. melanogaster* immune genes on a gene family level and found a high average conservation of 84.86% but a large difference for antimicrobial peptides with only one *O. bicornis* homologue compared to 22 *D. melanogaster* genes.

**Figure 2.**
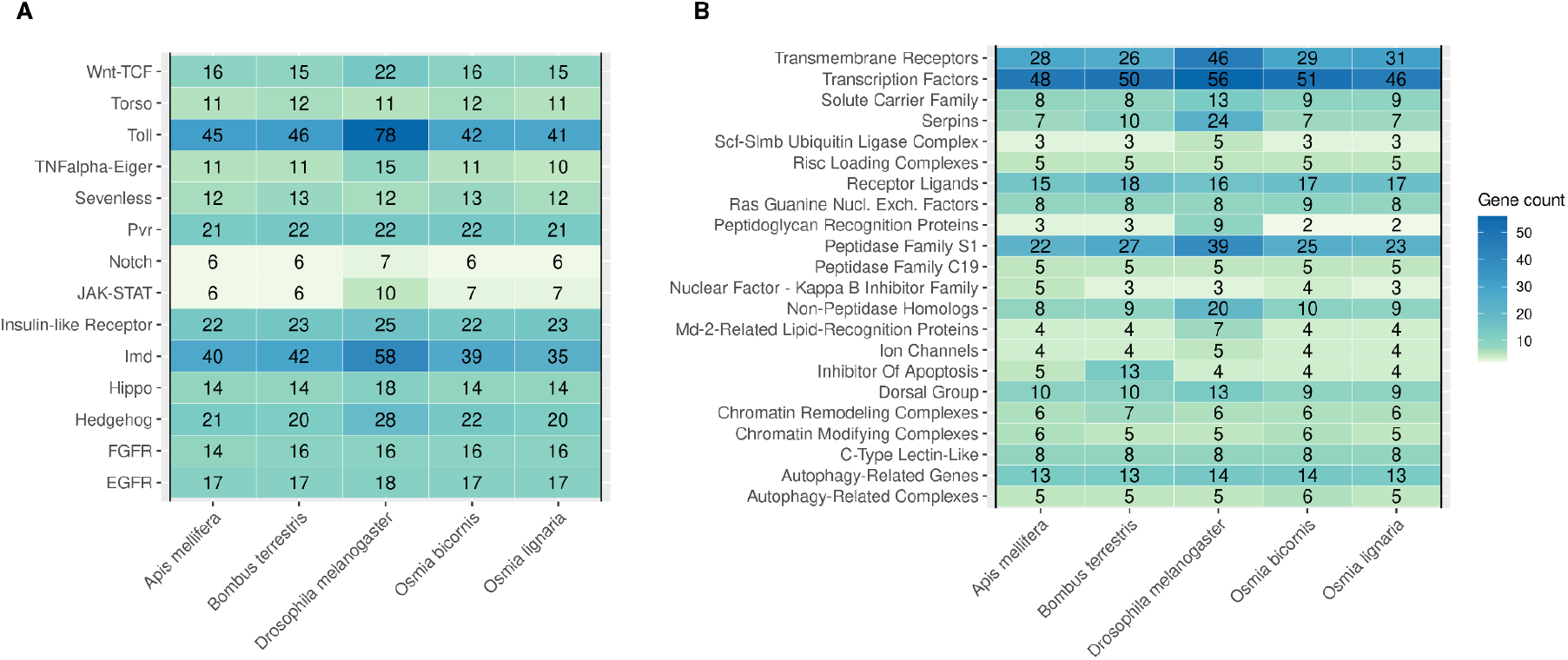
Conservation of immune genes and pathways in *Osmia bicornis*. Heatmaps depicting gene counts of homologues of putative *O. bicornis* immune genes for **(A)** molecular signaling pathways and **(B)** non-pathway gene families in closely related hymenopterans (*Apis mellifera, Bombus terrestris, Osmia lignaria*) and the more distantly related *Drosophila melanogaster*. For each species the number of homologues are shown.

### Sex-biased differential expression of immune genes

To provide functional information on the expression of putative immune genes in *O. bicornis*, we compared whole-bodied transcriptomes of male (n=3) and female adults (n=4). We identified 4,128 genes (34.99% of total gene count, n=11,799) as significantly differentially expressed (Likelihood-ratio test, BH-adjusted *p*<0.05) between the sexes, of which, 2,087 and 2,041 had female- and male-biased expression, respectively (Supplemental File S3). Among the differentially expressed genes, we found a significant enrichment or depletion (Kolmogorov-Smirnov test, *p*<0.05) of 125 biological process-associated GO terms, 47 cellular component GO terms and 29 molecular function GO terms with “cytoplasmic translation”, “cytosolic large ribosomal subunit” and “structural constituent of ribosome” as the most enriched terms for each of the three ontologies, respectively (Supplemental File S2). We quantified expression of 520 putative immune genes (96.11% of total immune genes, n=541) in both sexes, of which 222 were differentially expressed (42.69% of total DEGs) which was significantly more than expected by chance (Fisher’s Exact test, *p*=0.017). These differentially expressed genes were nearly equally shared between the sexes with slightly more genes (n=118) showing male-biased rather than female-biased expression (n=104), a pattern which did not significantly differ from expectation (Fisher’s Exact test, *p*=0.554, Supplemental File S1). Among the differentially expressed immune genes, we found enrichment or depletion for 28 biological process GO terms, five cellular component GO terms and five molecular function GO terms with “RNA localization”, “intracellular non-membrane-bounded organelle” and “RNA binding” as top enriched terms of each of the ontologies, respectively (Figure 3B, Supplemental File S2). We also analysed the *Osmia*-specific genes for signatures of differential expression and found 15 genes that significantly differed in their expression between the sexes (Supplemental File S1).

**Figure 3.**
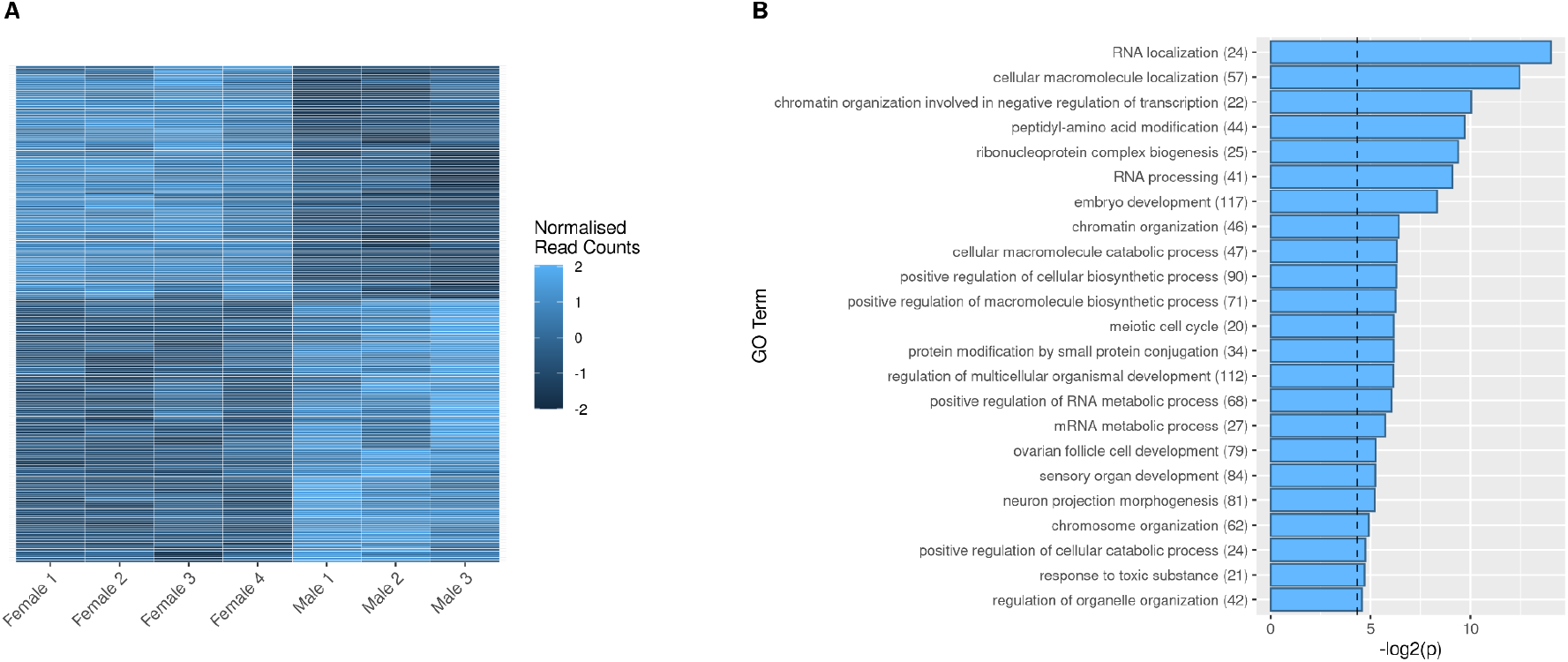
Differential expression of putative *Osmia bicornis* immune genes between males and females. **(A)** Heatmap showing normalised (variance stabilisation transformed) read counts per sample for significantly differentially expressed (BH adjusted *p* < 0.05) immune genes between males and females. **(B)** Functional enrichment of biological process GO terms of significantly differentially expressed putative *O. bicornis* immune genes. The x axis depicts negative log-transformed *p* values with the dashed vertical line corresponding to a *p* value confidence threshold of 0.05. Each line on the y axis corresponds to a GO term description with the numbers in brackets indicating the number of genes annotated with that specific term in the GO term database.

### Sex-biased alternative splicing of immune genes

We also performed alternative splicing analysis and identified 1,019 genes (8.64% of total gene count, n=11,799) significantly differentially spliced (Likelihood-ratio test, FDR < 0.05) between the sexes (Supplemental File S4). These sex-biased genes were significantly enriched or depleted (Kolmogorov-Smirnov test, *p*<0.05) for 107 biological process GO terms, 24 cellular component GO terms and 18 molecular function GO terms with “sarcomere organisation”, “Z disc” and “actin binding” as the top enriched terms for each ontology, respectively, as well as “immune system process”, significantly enriched among the biological process GO terms (Supplemental File S2). We identified 71 putative immune genes as differentially spliced (13.12%) between the sexes, which is significantly more than expected (Fisher’s exact test; *p*=0.002). In addition, we found differences between the sexes in the frequency of different splicing events with retained intron events significantly more common in females than in males (Fisher’s exact test, BH-adjusted *p*=0.036). For other splicing events, we found a borderline significant male-bias for alternative 3’ splice sites (Fisher’s exact test, BH-adjusted *p*=0.051) while we found no significant differences (BH-adjusted *p*>0.05) for skipped exons, alternative 5’ splice sites and mutually exclusive exons.

### Immune gene expression changes in response to pesticide exposure

For the neonicotinoid exposure analysis we identified 617 genes, including 42 putative immune genes, significantly differentially expressed (Wald-test, BH-adjusted *p*<0.05) in the group of thiacloprid-exposed females compared to the untreated control group (Figure 4C, Supplemental File S3), with a significantly unequal partitioning of 436 genes up-regulated and 181 genes down-regulated in the thiacloprid-exposed group (Binomial-test, *p*=9.41e-22). Comparing imidacloprid-exposed females with untreated females, we found 127 genes significantly differentially expressed (Wald-test, BH-adjusted *p*<0.05), including seven immune genes (Figure 4A). We also found a significantly unequal partitioning of 89 genes up-regulated and 38 genes down-regulated in the imidacloprid-exposed group (Binomial-test, *p*=6.97e-5). We found more differentially expressed immune genes than expected by chance in the thiacloprid-exposed group (n=42 immune genes; Fisher’s exact test, *p*=0.014) but not in the imidacloprid-exposed group (n=seven immune genes; Fisher’s exact test, *p*=0.515). Five putative immune genes (LOC114878095, LOC114878683, LOC114881181, LOC114874985, LOC114872156) had increased transcript expression both in response to thiacloprid and imidacloprid with no significant difference between the log2 fold change values for these five genes between the two pesticide treatment groups (Welch’s t-test, *p*=0.76, Supplemental Files S1, S3). In terms of functional enrichment of significantly differentially expressed immune genes, we found 45 biological process-associated GO terms enriched or depleted for the thiacloprid-exposed group, as well as five cellular component and eight molecular function terms with “regulation of hemocyte proliferation”, “integral component of plasma membrane” and “signaling receptor activity” as top enriched terms for each ontology, respectively (Figure 4D). For the imidacloprid-exposed group, we found 30 biological process GO terms enriched or depleted among the differentially expressed immune genes as well as four cellular component terms and seven molecular function terms with “transmembrane receptor protein tyrosine kinase signaling pathway”, “integral component of plasma membrane” and “signaling receptor activity” as top terms of each of the ontologies, respectively (Figure 4B, Supplemental File S2). Among the *Osmia*-specific genes we identified three genes differentially expressed between the thiacloprid-exposed group and control group but no differentially expressed genes between the imidacloprid-exposed group and the control group (Supplemental File S1).

**Figure 4.**
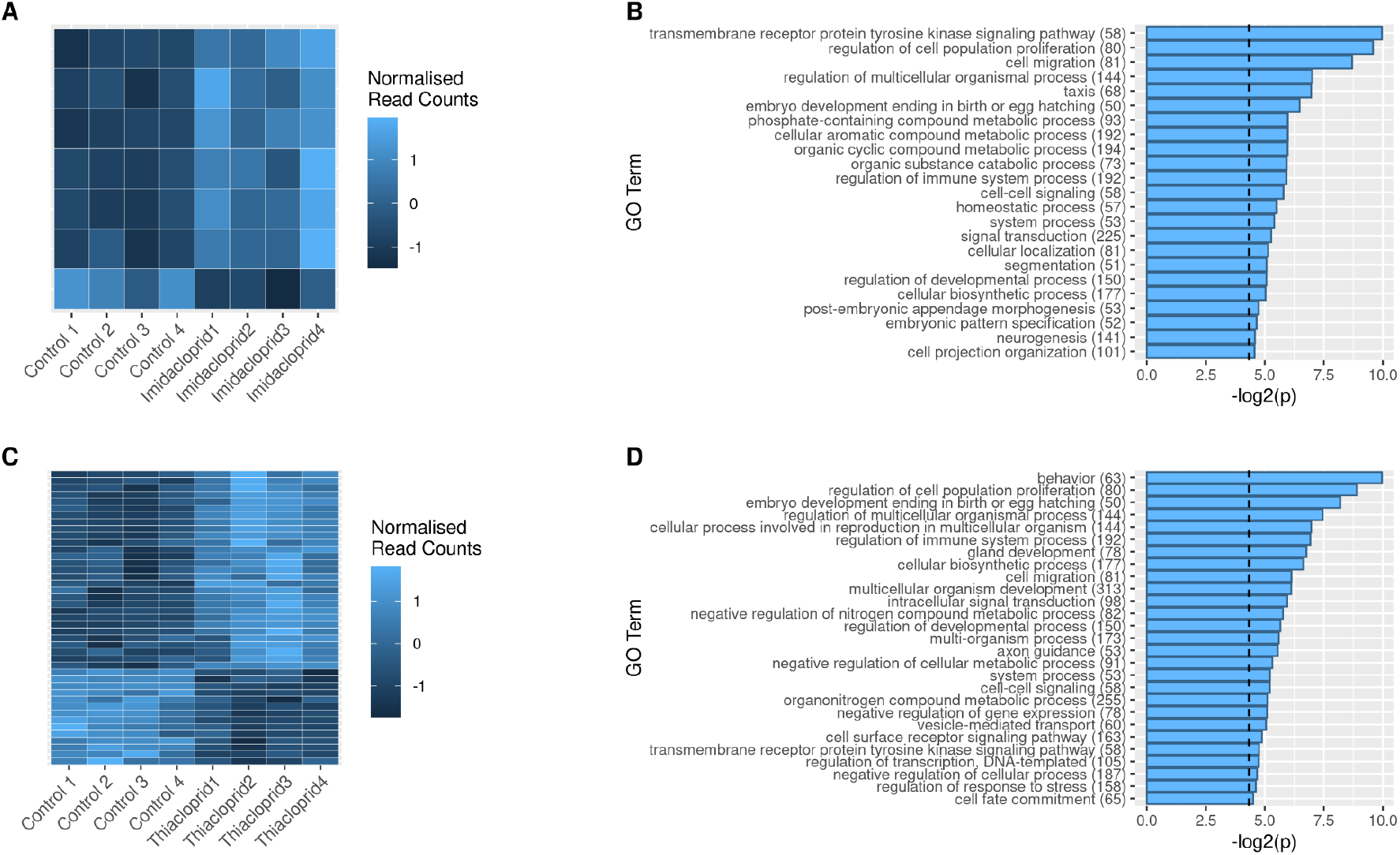
Differential expression of putative *Osmia bicornis* immune genes between pesticide-exposed and control individuals. **(A**,**C)** Heatmap showing normalised (variance stabilisation transformed) read counts per sample for significantly differentially expressed (BH adjusted *p* < 0.05) immune genes between control individuals and **(A)** imidacloprid-treated or **(C)** thiacloprid-treated individuals. **(B**,**D)** Functional enrichment of biological process GO terms of significantly differentially expressed putative *O. bicornis* immune genes between control individuals and **(B)** imidacloprid-treated individuals or **(D)** thiacloprid-treated individuals. The x axis depicts negative log-transformed *p* values with the dashed vertical line corresponding to a *p* value confidence threshold of 0.05. Each line on the y axis corresponds to a GO term description with the numbers in brackets indicating the number of genes annotated with that specific term in the GO term database.

For alternative splicing, we found 142 differentially spliced genes (Likelihood-ratio test, FDR<0.05) for the thiacloprid-exposed group, including six putative immune genes, while 74 genes were differentially spliced for the imidacloprid-exposed group, including four putative immune genes. One immune gene (LOC114880106) was differentially spliced in response to both pesticides (Supplemental File S4).

## Discussion

The insect immune system represents an important barrier against infections and disease and thus provides a physiological function vital to an individual’s success. While our understanding of the genomic and molecular bases of immunity in the Hymenoptera has largely been informed by studies on social bees, for other species, especially solitary bees, such information is limited. Here, we performed a comparative genomic analysis to characterise genes with potential immune functions in the genome of the red mason bee, *O. bicornis*. Using a homology-based approach, we identify an immune gene repertoire enlarged beyond the canonical immune genes previously described in other hymenopteran genomes. We find extensive differences in immune gene expression between the sexes, both in terms of expression amplitude and splicing, highlighting intrinsic regulatory differences in the molecular basis of immunity between males and females. Lastly, we find immune-related genes differentially expressed in response to neonicotinoid exposure with greater expression differences in bees exposed to thiacloprid than those exposed to imidacloprid demonstrating differences in how the molecular phenotype responds to different neonicotinoid subclasses and how each may influence immune expression.

The insect immune system consists of an innate immune response with the ability to detect and remove a diverse range of pathogenic entities (Beckage, 2008). In addition to behavioural, physical and chemical defences, it is an important barrier to infection and disease development. The earliest genomic studies on the Hymenoptera documented a reduction in canonical immune genes in comparison to other insect orders, most notably in comparison to members of the Diptera (Evans *et al.*, 2006; Bonasio *et al.*, 2010; Werren *et al.*, 2010; Barribeau *et al.*, 2015). Reasons for this reduction ranged from the technical (e.g., fragmented genome assembly, missing or truncated gene models) to the biological level (e.g., novel immune genes and pathways ((Albert *et al.*, 2011; Dong *et al.*, 2020) or relaxed selection acting on canonical immune genes due to social immunity (Harpur and Zayed, 2013)). Here we performed one of the first investigations of the immune gene repertoire of a solitary bee species.

Our initial approach for the detection of putative immune genes was based on homology with genes annotated with roles in immune system function based on Gene Ontology in the model organism, *D. melanogaster*, where many such genes have been experimentally validated with roles in immune function. As many immune genes have been previously shown to evolve under strong episodic positive selection (Jiggins and Kim, 2007; Viljakainen, 2015; Shultz and Sackton, 2019), which can result in divergence beyond detection through homology searches or the appearance of novel lineage-specific immune genes, we also investigated homologues in *O. bicornis* of canonical immune genes from the earth bumblebee, *B. terrestris*, a closely related social insect with an annual life-cycle. As the majority of canonical immune genes identified in *B. terrestris* were generated based on homology searches with other insect genomes, including *D. melanogaster* (Barribeau *et al.*, 2015), we would have expected canonical immune genes in both species to be annotated with immune process GO terms and therefore, we would have expected to see a high overlap in immune gene sets. Surprisingly, we found a weak overlap between *Osmia* homologues identified using both approaches, with only a third of *B. terrestris* canonical immune genes identified also through homology to *D. melanogaster* immune genes and thus annotated with immune system GO terms. The other two thirds were annotated with GO terms associated only more indirectly with immunity, such as “response to toxic substance”, “RNA interference” and “autophagy”. The lack of annotation of immune GO terms for two thirds of the described canonical immune genes in *B. terrestris* may result in the underreporting of immunological changes for genomic studies reliant on GO term based analyses.

To provide better support for conserved function for putative immune homologues in *Osmia*, we further assessed *O. bicornis* immune homologues identified through the *D. melanogaster* comparison, which lacked a described homologue in *B. terrestris* canonical immune genes. If such homologues were spurious or low quality matches, we predicted such protein homologues may have lower percentage sequence identity, greater differences in sequence length, as well as variation or lack of structural features, such as abundance and diversity of functional domains compared to canonical immune genes. However, for the majority of homologous pairs, we found the same or greater percentage sequence identity as canonical immune genes, as well as the presence and conservation of functional domains, suggesting that potential immune functions may be conserved and perform similar roles in bees. Identified through homology with *D. melanogaster* immune genes only, we found homologs of many immune relevant genes like defensins, hemocytin and sickie known to be important to the immune system across different insect species (Hoffmann and Hetru, 1992; De Gregorio *et al.*, 2002; Lavine and Strand, 2002; Arai *et al.*, 2013; Ni *et al.*, 2020). Through homology with *B. terrestris* immune genes only, however, we found homologues of immune genes known to be involved in the insect immune system, like mucins, galectin-like proteins and superoxide-dismutases (Pace and Baum, 2002; Korayem *et al.*, 2004; Colinet *et al.*, 2011; Rao *et al.*, 2016), demonstrating the importance of using more than one species to infer undescribed gene sets via homology. While future experimental studies on their function will elucidate potential roles in immunity, if any, for these additional candidate genes, their high number could also speak to the ever-increasing completeness of functional annotation in insect model organisms, like *D. melanogaster*.

Among the 541 putative immune genes of *O. bicornis*, we did not find any major patterns of immune gene family expansions or contractions in comparison with that of *B. terrestris* and *A. mellifera*, suggesting that there were no large-scale recent duplications or losses of genes that could be imperative to the functioning of the immune system of bees. However, we did find slight reductions in the number of genes involved in two major immune signaling pathways, Imd and Toll, when we compared *O. bicornis* and *O. lignaria* to *A. mellifera* and *B. terrestris*. Imd and Toll are both involved in the induction of antimicrobial peptides (AMPs) (De Gregorio *et al.*, 2002), thus, fundamental for insect survival in response to pathogen challenge. Related to this, the biggest difference in immune gene families between *O. bicornis* and *D. melanogaster* was for gene copies of AMPs. This can be expected as *Drosophila* have evolved a number of AMP gene families, such as the cecropins, diptericins and attacins (Imler and Bulet, 2005), which have not been identified in other hymenopteran genomes (Evans *et al.*, 2006; Barribeau *et al.*, 2015). Consequently, we did identify the presence of defensin, an evolutionary conserved AMP that possesses antibacterial properties (Hoffmann and Hetru, 1992), as well as the hymenopteran-specific AMP, hymenoptaecin (Casteels *et al.*, 1993). However, we did not detect a copy of abaecin, a bacterial-inducible AMP described in honeybees (Casteels *et al.*, 1990) and bumblebees (Rees, Moniatte and Bulet, 1997). The conserved lack of abaecin, as well as components of the Imd and Toll signaling pathways, in the genome assemblies of two *Osmia* species, which were sequenced and assembled independently, suggests that such missing genes may be true biological signals rather than technical artefacts. At present the evolutionary consequences of such potential losses, if any, are unknown, but it suggests at least that differences in the molecular structure of the immune system do indeed exist for these two solitary bee species compared to these other social bee species.

For species that sexually reproduce, the genome codes for distinct sexes that can differ dramatically in behaviour, morphology and physiology (Parsch and Ellegren, 2013), including immunity (Klein and Flanagan, 2016). As immunity can be both energetically costly to maintain and activate (Rolff and Siva-Jothy, 2003), it can result in metabolic trade-offs with other processes, such as reproduction (Schwenke, Lazzaro and Wolfner, 2016). Sexes can also differ in immune investment, which may be reflected in differences in gene expression. Here we found 222 putative immune genes to be differentially expressed between the sexes with the strongest functional enrichment for terms related to localization and breakdown of RNA and general macromolecules. This suggests that the sexes may differ in important housekeeping roles, such as RNA and protein turnover, but also how they respond to challenging macromolecules that need to be localized and decomposed, such as RNA-viruses and toxins. The case for differences in antiviral defence is further highlighted by genes, such as endonuclease Dcr-1 or defensin, which have roles in virus recognition and degradation (Galiana-Arnoux *et al.*, 2006; Brutscher, Daughenbaugh and Flenniken, 2015), being differentially expressed between the sexes. We found more putative immune genes than expected by chance to be differentially expressed or differentially spliced between the sexes suggesting that the molecular differences between the sexes are particularly pronounced when it comes to immune system processes. We also looked at the expression differences between sexes in general and found the most striking functional enrichment of differentially expressed genes to be related to translational and cell division processes, underlining the generality of molecular differences between the sexes while the most striking enrichment of alternatively spliced genes is mostly related to muscle activity and regulation of muscle excitation which might reflect the different life strategies of male and female mason bees where male bees emerge prior to female bees or thermoregulatory differences.

Pesticides, including neonicotinoids, act as agonists of the nicotine acetylcholine receptors, resulting in disruption of the neuronal cholinergic signal transduction and excitation of neuronal triggers culminating in paralysis and death (Matsuda *et al.*, 2001). The efficacy of the mode of action of neonicotinoids has led to their increased popularity in modern agriculture practices yet recent studies have highlighted the negative impact sublethal and lethal doses can have on pollinator health (‘Neonicotinoids, bee disorders and the sustainability of pollinator services’, 2013), including immune function. Neonicotinoids, such as clothianidin and imidacloprid, have been identified in exposed honeybees to negatively modulate the NF-kappaB signaling pathway and affect the ability to mount effective antiviral defences (Di Prisco *et al.*, 2013). Other studies on neonicotinoids have provided additional evidence of the indirect or direct effects of these neurotoxins on immune function or expression (Brandt *et al.*, 2017, 2020). Here we found changes in gene expression in response to two classes of neonicotinoids with thiacloprid exposure affecting the expression of more genes, including immune genes, than imidacloprid. This is in line with other studies that have looked at thiacloprid exposed red mason bees and observed impairment in immunity (Brandt *et al.*, 2020) or larval development (Claus *et al.*, 2021). For both neonicotinoids, we find significantly more genes up-regulated in response to pesticide exposure than expected, suggesting that overall the exposure to pesticides results in an active response of heightened gene expression as opposed to a mere passive shift in gene expression. Focusing on immune system processes, we find more immune genes differentially expressed than expected only in thiacloprid-treated individuals, suggesting that thiacloprid elicits a stronger immune response than imidacloprid. Interestingly, of the seven differentially expressed genes with elevated expression in imidacloprid-exposed bees, five genes were also significantly up-regulated in the thiacloprid-exposed bees which could possibly point to a common set of immune genes that are up-regulated in response to neonicotinoid exposure. In terms of functional enrichment of the differentially expressed immune genes, both pesticide-treated groups share similar terms, related to signaling of transmembrane receptors, suggesting that the immune genes play a role in signaling in response to pesticide exposure. In addition, in the thiacloprid-treated group the term “regulation of hemocyte proliferation” is the most significantly enriched term. Haemocytes fulfill an important role in the ingestion and break-down of foreign cells and substances in the insect immune system (Lavine and Strand, 2002). Brandt *et al*. found a reduction in haemocyte density in red mason bees after exposure to thiacloprid, which is in line with our finding of haemocyte proliferation being differentially regulated between thiacloprid treated and untreated individuals, suggesting that the effect of pesticide exposure on haemocyte function may extend down to the molecular level.

## Conclusions

The red mason bee, *O. bicornis*, is a commercially and ecologically relevant solitary bee species, whose immune system has not been well-studied yet, albeit being integral to its future chances of survival when faced with increasing environmental challenges. Here, we utilised a comparative genomic approach to propose a set of genes as part of the immune gene repertoire of *O. bicornis*, and used RNA-seq data to show that the expression and regulation of these putative immune genes differs markedly between sexes and responds with heightened expression to treatment with two neonicotinoid pesticides. Additionally, our findings provide support for the application of a combined approach to inference of gene families, using homology information of more than one species of reference. Future studies on *O. bicornis* immunity will benefit from tissue-specific profiling, as well as tracking gene expression changes in response to different immune challenges. Similarly the application of population genomics will provide important insights into the recent selection pressures acting on immune genes of mason bees. Collectively, our study provides novel insights into the immune system of an important, yet still understudied solitary bee species and identifies a candidate repertoire of immune genes for future research on the immune system of the red mason bee.

## Experimental Procedures

### Identification of putative immune genes in the red mason bee

To infer homologues for *O*.*bicornis* genes that are in other insect species, we ran OrthoFinder [v.2.5.2](Emms and Kelly, 2019) with proteomes of 21 species, comprising 11 bee species from three families (Family Megachilidae: *Osmia bicornis*, O*smia lignaria*, and *Megachile rotundata*; Family Apidae: *Apis mellifera, Ceratina calcarata, Eufriesea mexicana, Habropoda laboriosa* and *Bombus terrestris*; and Family Halicitidae: *Megalopta genalis, Nomia melanderi* and *Dufourea novaeangliae*), as well as 10 non-bee insects (*Drosophila melanogaster, Aedes aegypti, Anopheles gambiae, Bombyx mori, Tribolium castaneum, Acyrthosiphon pisum, Nasonia vitripennis, Solenopsis invicta, Polistes dominula* and *Vespa mandarinia*). All proteomes were obtained from the National Center for Biotechnology Information (NCBI) Reference Sequence (RefSeq) database. We ran OrthoFinder using the default parameters with the inferred species trees forming a consensus with the known phylogeny. Given that model organisms, such as *D. melanogaster*, contain the most comprehensive functional annotation of genes with immune function or potential, we examined the *O. bicornis* predicted proteome for putative homologues of *D. melanogaster* immune genes. To obtain *D. melanogaster* immune-responsive genes, we queried the FlyBase on the 9th of July, 2021 for any gene associated with the GO term “immune system process”, the highest order Gene Ontology term associated with the immune system, and inferred the *O. bicornis* homologues via the homologues table generated earlier. As an additional approach, to identify immune genes that may be lineage-specific within the Hymenoptera or too divergent between *O. bicornis* and *D. melanogaster* given their evolutionary distance, we examined the presence of *O. bicornis* homologues from comparison with canonical immune genes characterised in the earth bumblebee, *Bombus terrestris*. The canonical *B. terrestris* immune genes were directly obtained from the most recent earth bumblebee genome papers (Barribeau *et al.*, 2015; Sadd *et al.*, 2015). An additional reason for the inclusion of this social bee species is that it has a curated homologue list with *D. melanogaster*, available through the Ensembl Metazoa database, which provided the ability to compare the orthogroups and homologous pairs generated by OrthoFinder with orthogroups and pairs independently generated by Ensembl, which incorporates an additional information on synteny and gene order conservation for the identification of putative homologues between two species. This approach identified a high overlap (86.85% of Ensembl pairs correctly identified by OrthoFinder) between the pairs generated by both analyses providing additional confidence in the orthogroups generated by OrthoFinder. Homologous genes from *D. melanogaster* and *B. terrestris* were obtained by first translating the *O*.*bicornis* gene-IDs to protein-IDs via the annotation column in the RefSeq gene feature file (GFF) and then using the homology information from OrthoFinder to translate *O. bicornis* protein-IDs to *D. melanogaster* and *B. terrestris* protein-IDs respectively and further translating the protein-IDs to the species-specific gene-IDs, yielding a many-to-many homologue table of *O. bicornis* gene-ID’s to *D. melanogaster* and *B. terrestris* gene-ID’s, respectively. To infer for each gene family how many homologous genes exist for the set of putative *O. bicornis* immune genes in *A. mellifera, B. terrestris, D*.*melanogaster* and *O. lignaria*, we derived annotation of *D. melanogaster* genes with gene families from Flybase on the 02nd of June, 2021 and annotated the immune gene homologues in each species with the according gene family description. We then summarised this data by counting the number of genes in each species and for each gene family.

For *O. bicornis* genes that shared homology with *D. melanogaster* immune-responsive genes but did not overlap with known canonical immune genes in *B. terrestris*, we further examined homology based on the following criteria: 1) similarity of predicted protein length between homologous pairs; 2) high percentage of protein sequence identity between homologous pairs as inferred via Diamond searches performed by OrthoFinder; and 3) the number of shared functional protein domains between homologous as inferred via InterProScan [v5.52-86.0](Jones *et al.*, 2014). The prediction here is that if two homologous proteins shared similar protein length, high sequence identity and the same or similar number and types of functional protein domains, potential functional immune roles may be conserved.

In addition to identification of immune-related genes, we also inferred canonical immune genes from *B. terrestris* or *D. melanogaster* missing in *O. bicornis and O. lignaria*. For this, we parsed the output of OrthoFinder for orthogroups that carried immune-associated genes in *D. melanogaster* or *B. terrestris* but not in both *O. bicornis* and *O*.*lignaria*. Similarly, we inferred *Osmia*-specific genes by parsing orthogroups containing only *O. bicornis* and *O. lignaria* homologues, which were also absent in the other 19 species.

### Quality assessment, transcript abundance estimation and differential expression analysis of immune genes between sexes and pesticide-treated groups

To examine the functional expression of putative immune genes of *O. bicornis*, we obtained publicly available paired-end RNA-seq libraries for two analyses: a) sex-biased analysis containing males (n=3) and females (n=4); and b) pooled libraries of unexposed (“control”; n=4) females or those exposed to thiacloprid (n=4) or imidacloprid (n=4). All datasets were obtained from the NCBI (National Center for Biotechnology Information) Short Read Archive (SRA) database (BioProject: PRJNA285788; (Beadle *et al.*, 2019), Supplemental File S5). We performed data quality assessment based on per-sample quality evaluations using FastQC [v.0.11.9](Andrews, 2010) calculation of the proportions of reads mapping to the predicted transcriptome of the RefSeq *O. bicornis* reference genome assembly [Obicornis_v3; GCF_004153925.1] using Kallisto [v.0.46.1] (Bray *et al.*, 2016). We then combined and visualized the results for both tools and across all samples with MultiQC [v.1.7](Ewels *et al.*, 2016). Based on the results of the quality assessment we removed adapter sequences and filtered by quality (phred quality score >= 15) and length (minimum length >= 50 bp) using fastp [v.0.20.1](Chen *et al.*, 2018). For each sample, we then aligned the trimmed and filtered reads against the most recent chromosome-level genome assembly [iOsmBic2.1; GCA_907164935.1] using STAR [v.2.7.8a](Dobin *et al.*, 2013). As the chromosome-level assembly currently lacks annotations, we first transferred gene coordinates from the annotated reference assembly [Obicornis_v3; GCF_004153925.1] to the new assembly using Liftoff [v.1.6.1](Shumate and Salzberg, 2020). STAR was ran in two-pass-mode using the inferred splice junctions from the first run to improve the alignment of the second run and with parameter --quantMode GeneCounts used to generate gene level abundances of aligned reads (Supplemental File S6). The mean alignment rate across all samples was 94.09%. For the sex-biased analysis, we used DESeq2 [v.1.30.1](Love, Huber and Anders, 2014) to correct for library size and infer all differentially expressed genes (“DEG”, FDR < 0.05) between sexes with a likelihood-ratio-test (LRT; full model: sex; reduced model: intercept). For the pesticide-based analysis, we implemented pairwise Wald tests to determine log2 fold changes between each pesticide treatment and control, as well as quantify all differences in gene expression between pesticide treatments. For each analysis, we then parsed DEGs for putative immune genes. To determine if we identified more or less immune genes than would be expected, we performed Fisher’s exact tests for each analysis.

As a complementary approach to STAR, we also implemented a pseudoalignment-based differential gene expression analysis using Kallisto [v.0.46.1] as described above in the quality assessment section (mean mapping rate across samples 87.93%). Similar to the STAR-based analysis, we implemented the same statistical tests using DESeq2 for both the sex-biased and pesticide-based analysis, respectively. Out of the genes predicted to be differentially expressed using STAR, 3376 genes (81.78%, n = 4128) were also reported to be differentially expressed using Kallisto and inversely, 92.01% (n = 3669) of all genes predicted to be differentially expressed using Kallisto were also predicted to be differentially expressed using STAR.

### Splicing of immune genes between sexes and pesticide-treated groups

To determine differentially spliced genes between the sexes, as well as in response to pesticide exposure, we ran rMATS turbo [v.4.1.1](Shen *et al.*, 2014) using the STAR generated alignment files. Similar to our differential expression analyses, we performed two independent analyses: first, we compared males and females to identify significant sex-biased differences in splicing; second, we performed a pesticide-based analysis, comparing thiacloprid- and imidacloprid-exposed females against the control group, respectively. A specific feature of rMATS is that it outputs event-level instead of gene-level results, distinguishing between different event types such as exon-skipping, intron-retainment and 3’- or 5’-splicing and allowing multiple splicing events to be recognized per gene. We used this event-based information to investigate whether there were incidences of alternative splicing where a specific splicing event appears preferentially in one of the experimental groups over the other. To do this we compared the number of times a differential splicing event had higher inclusion levels in one group over the other against the total number of incidences of differential splicing for that event using a Binomial-test with the null hypothesis being that we would see higher inclusion levels equally often for both groups. We then concatenated and summarised the event-level results by grouping the events by gene-ID and summarising down to the lowest *p* value per gene, yielding a gene-level output table.. We then used this information to rank genes by *p* value and investigate functional enrichment of Gene Ontology (GO) terms in alternatively spliced genes (below).

### Gene Ontology term enrichment analysis

As most genes in the *O. bicornis* genome lack functional information, we assigned GO terms to genes from homologues found in *D. melanogaster*, which were obtained from Ensembl Metazoa via biomaRt [v.2.46.3](Durinck *et al.*, 2009; Kinsella *et al.*, 2011). We then ran functional enrichment analyses using topGO [v.2.42.0; (Alexa and Rahnenführer, 2009)] on the sets of immune genes derived via homology to A) *D. melanogaster* uniquely, B) *B. terrestris* uniquely and C) to both species. To do so we implemented the “classic” algorithm in combination with a Fisher test (node size=20) using all genes, each marked for presence or absence in the set of putative immune genes. We then tested for enrichment of the most differentially expressed genes between sexes and pesticide treatment groups. For this approach, we ranked genes by unadjusted raw *p* values to avoid edge-effects introduced by correction and performed a rank-based analysis (Kolmogorov-Smirnov test with “weighted01” algorithm and a node size=20). We also performed an immune-focused GO term enrichment analysis where we populated our GO term database only with genes annotated with immune-related GO terms.

## Data and Code Availability

Data used for the present analysis originated from datasets generated by (Beadle *et al.*, 2019). Raw sequences for the RNA-seq based analyses can be obtained from the NCBI Short Read Archive (BioProject PRJNA285788).

